# Role of SCAL1 in Modulating Oxidative Stress, Cancer Stemness, Apoptotic Resistance in Tumorigenic Differentiation of Cigarette Smoke-Exposed BEAS-2B Cells

**DOI:** 10.1101/2024.08.05.606632

**Authors:** Debmalya Sengupta, Souradeep Banerjee, Mainak Sengupta

**Affiliations:** Department of Genetics, University of Calcutta, Kolkata, INDIA Institutions where the work has been carried out: (i) Department of Genetics, University of Calcutta, Kolkata, INDIA

**Keywords:** Lung cancer, Cigarette smoke extract, SCAL1, BEAS-2B cells, Cancer stem cell markers

## Abstract

The smoke and cancer-associated lncRNA 1 (SCAL1) is an emergent biomarker in lung cancer. However, the precise role of SCAL1 as a mediator of tobacco smoke-induced lung carcinogenesis remains unclear. BEAS-2B cells were cultured and exposed to 20% cigarette smoke extract (CSE), followed by quantification of SCAL1. We evaluated the impact of SCAL1 on cell viability, ROS mitigation, cancer stemness, tumorigenic differentiation, cellular invasiveness, and apoptosis for different CSE incubation time points through SCAL1 expressional modulation using SCAL1-specific siRNAs and scrambled controls. We observed an upregulation of SCAL1 in cells exposed to CSE for 2, 4, and 6 hours, with the highest expression observed at 6 hours (p<0.001). Exposure of BEAS-2B cells to CSE showed the formation of focal adhesions and stress fibers resembling tunneling nanotubes. Intracellular ROS levels significantly increased upon CSE exposure compared to control cells (p<0.05). We found increased levels of anti-apoptotic and cancer stem (CSC) cell markers like BCL2, ALDH1A1, CD133, CD44, and TCTP and decreased levels of TP53 in CSE-exposed cells. Knockdown of SCAL1 using siRNA transfection reversed these effects at all time points. Additionally, we observed a significant decrease in the number of spheroid colonies in siSCAL1 (+) cells compared to siSCAL1 (-) cells (p<0.01) exposed to CSE. SCAL1 is pivotal in mediating cellular responses to cigarette smoke, leading to tumorigenic differentiation of BEAS-2B cells. Understanding the mechanisms could provide valuable insights into lung cancer pathogenesis and therapeutic approaches.

## Introduction

Lung cancer ranks second in incidence and first in cancer-related mortalities worldwide [1], with tobacco smoking as the primary etiological factor [2]. Tobacco smoke induces genetic and epigenetic reprogramming of airway cells [3], cellular dedifferentiation [3, 4], and phenotypic plasticity [4–6], leading to severe lung malignancies.

Long non-coding RNAs (lncRNAs) are non-protein coding transcripts that are >200 nucleotides [7] and recruit transcription factors (TFs) to regulate gene expression or interact with microRNAs (miRNAs) to modulate the mRNA stability [8]. LncRNAs are potential biomarkers for cancer diagnosis, prognosis, and therapy [9], involved in epigenetic regulation in pathological conditions [8, 9] such as tumorigenesis and its progression. The smoke and cancer-associated lncRNA 1 (SCAL1) is an 890bp lncRNA, also known as lung cancer-associated transcript 1 (SCAL1), coded by its gene located at 5q14.3, was first detected in the airway epithelium of smokers [9, 10] found to be elevated in lung cancer cell lines and thus identified as a tumor-related lncRNA induced by cigarette smoke. SCAL1 is overexpressed in NSCLC tissues with advanced TNM (Tumour, Node, Metastasis) stage, lymph node metastasis, and poor prognosis [11] compared to normal tissues.

Recent investigations have shown the role of SCAL1 in cisplatin resistance in ovarian and colorectal cancer [12]. Cigarette smoke exposure is known to upregulate SCAL1 through the NFE2L2/KEAP1 pathway that promotes cytoprotection and mitigation of smoke-induced ROS in lung cancer cell lines [10]. Another study showed that siRNA-mediated knockdown of SCAL1 arrests the cell cycle at the G_0_/G_1_ stage and decreases both the binding of EZH2 and H3K27me3 across the p21 and p57 promoters, which enhances the expression of p21 and p57 in NSCLC [11]. SCAL1 exerts an oncogenic function in cervical cancer by binding to miR-181a, showing its miRNA-mediated gene regulation [13]. SCAL1 binds to UBA52 to induce its degradation into ubiquitin molecules, which then regulate the stability of MDM2, an inhibitor of the tumor suppressor p53 [14]. SCAL1 inhibits KISS1 expression, promoting migration and invasion of prostate cancer cells. [15]. Additionally, the ribosomal protein RPL40, cleaved from UBA52, interacts with MDM2 and is involved in the RP-MDM2-p53 pathway, further linking SCAL1 to MDM2 regulation in prostate cancer [14]. SCAL1 inhibits Nucleolin (NCL) function via G-quadruplex, serving as a potential biomarker and therapeutic target for colorectal cancer (CRC) [16]. A recent study revealed SCAL1 increases cancer stemness by competitive binding of miR-5582-3p with TCF7L2 in breast cancer survivors [17]. SCAL1 functions as a molecular sponge for miR-539, mediating the effects of SCAL1 on pancreatic ductal adenocarcinoma (PDAC) cell proliferation, cell cycle progression, and motility [18]. The role of lncRNA in CSC maintenance and the pathway to block CSC regulation has been the primary focus of modern cancer research [19, 20]The available literature suggests that the mechanistic impacts of SCAL1 are evaluated in cancer cells and/or tissues. However, the precise role of SCAL1 in cytoprotection, tumorigenic differentiation, generation of cancer stem cell-like properties, and resistance to apoptosis in otherwise normal healthy human bronchial epithelia on CSE exposure is still elusive. It is imperative to assess the development of the tumor hallmarks in normal healthy human bronchial epithelial cells to understand tobacco smoke-dependent mechanisms of lung carcinogenesis.

The study, for the first time, aims to delineate the role of SCAL1 in ROS mitigation and cytoprotection, cell proliferation, apoptosis, generation, and maintenance of CSCs, followed by tumorigenic differentiation on CSE exposure to immortalized human primary bronchial epithelial (BEAS-2B) cells *in vitro*. The mechanism that induces CSC could be the dedifferentiation of epithelial cells when exposed to carcinogens in tobacco smoke metabolites. These metabolites generate significant ROS, imposing oxidative stress on the cells. The cells could acquire the ability to undergo transformation and adapt to the increased levels of reactive oxygen species (ROS), which could lead to the development of tumors. The proposed schema of experiments is illustrated in the following **Fig. 1**.

**Figure 1.**
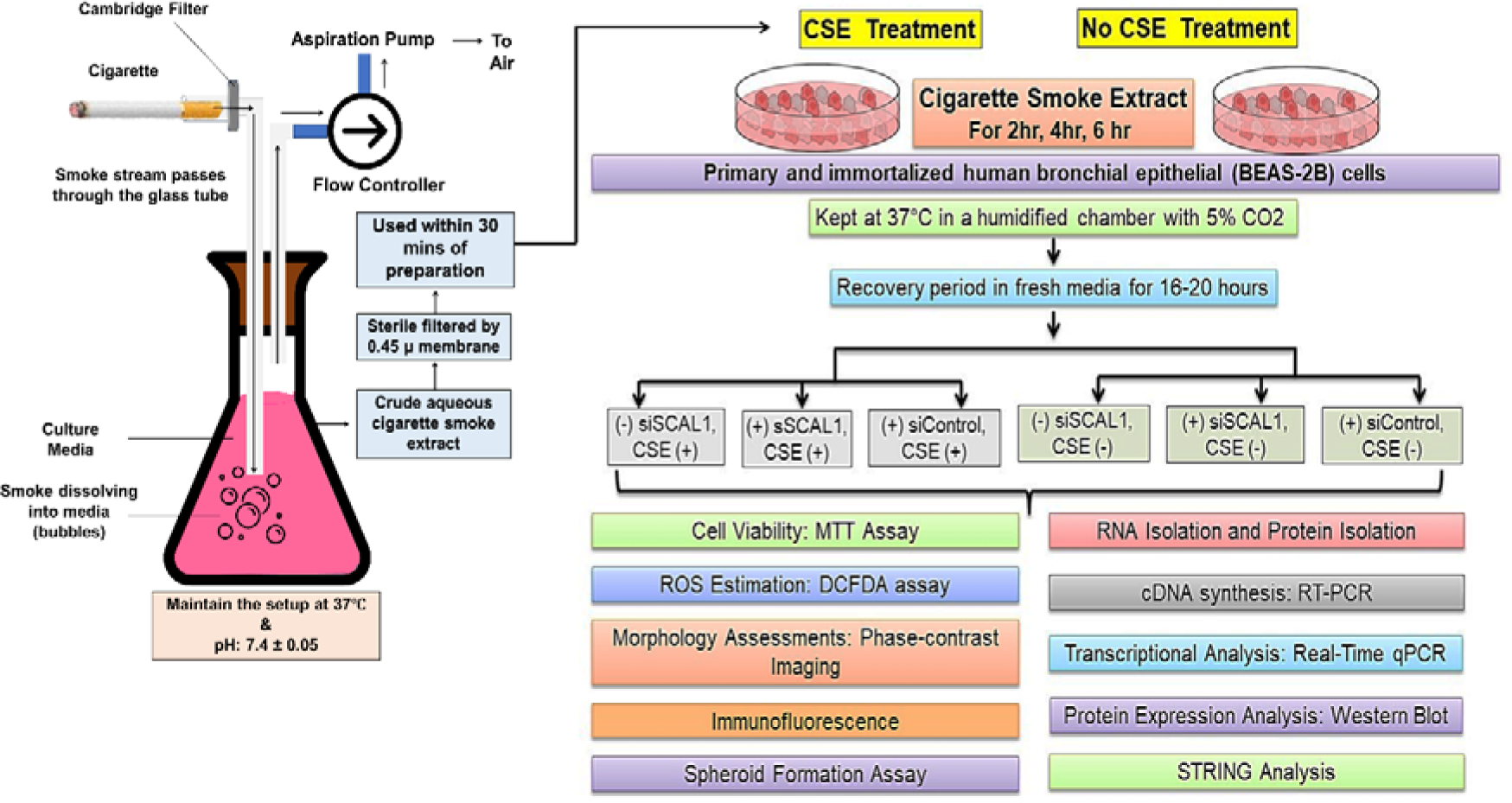
Schema of the study.

## Results

We have compared the outcomes for siSCAL1 (-), CSE (-) BEAS-2B cells with siSCAL1 (-), CSE (+) BEAS-2B cells for 2, 4, and 6 hours of incubation in 20% CSE. We have also compared the outcomes for siSCAL1 (-), CSE (+) BEAS-2B cells with siSCAL1 (+), CSE (+) BEAS-2B cells matched with the hours of incubation in 20% CSE and siSCAL1 (-), CSE (-) BEAS-2B cells with siSCAL1 (+), CSE (-) BEAS-2B cells. Furthermore, we have also compared the outcomes of siControl (+) BEAS-2B cells with all the groups matched with 20% CSE exposure and the hours of CSE exposure. The details of the outcomes for all the cases are delineated below. The siControl (+) cells showed similar outcomes as that of siSCAL1 (-) cells.

### Overexpression of SCAL1 in BEAS-2B cells under cigarette smoke exposure

Gene expression of SCAL1 increased significantly (*p*<0.05) after exposure to 20% CSE for 2, 4, and 6 hours (4.71-fold, 7.64-fold, and 13.25-fold, respectively) compared to the unexposed group. Silencing SCAL1 with siSCAL1 showed statistically significant downregulation of SCAL1 expression compared to scrambled siControl (**Fig. 2[A]**).

**Figure 2.**
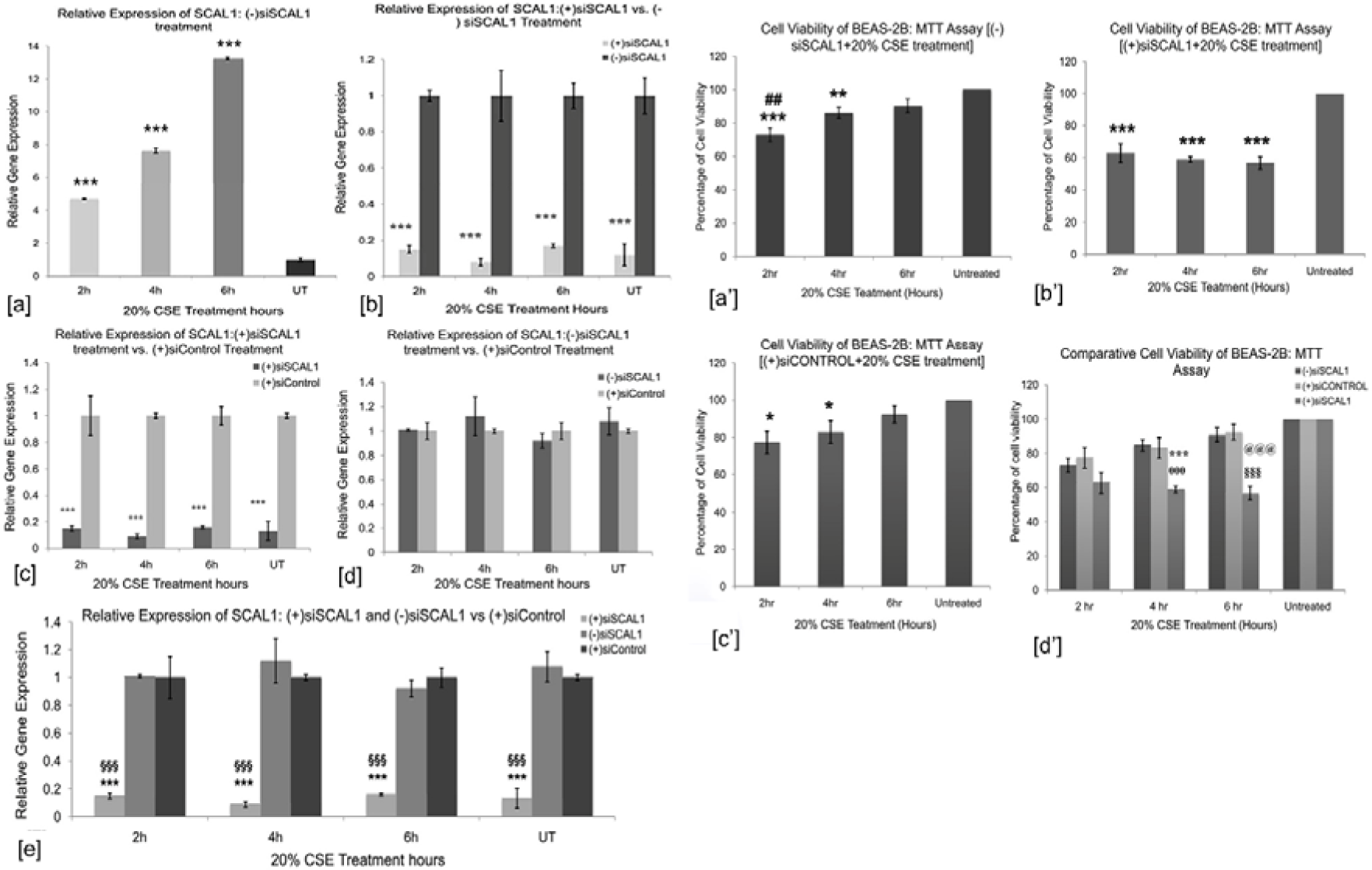
[A] Expression of SCAL1 by cigarette smoke extract (CSE) exposure. Primary immortalized bronchial epithelial (BEAS-2B) cells were exposed to 20% CSE for 2 hours, 4 hours, and 6 hours followed by a 16-20 hour recovery, [a] a statistically significant upregulation of SCAL1 expression in siSCAL1 untreated, [b] (-) siSCAL1, (+) 20% CSE *vs.* (+) siSCAL1, (+) 20% CSE, [c] (+) siSCAL1, (+) 20% CSE vs. (+) siControl, (+) 20% CSE, [d] (-) siSCAL1, (+) 20% CSE vs. (+) siControl, (+) 20% CSE, each of them has 20% CSE untreated as Control set Combined representation of (+) siSCAL1 and (-) siSCAL1 vs. (+) siControl. *Error bars represent SEM. P-values estimated by post hoc t-test (Bonferroni method). P<0.05*, 0.01**, 0.001*** (comparison between (-) siSCAL1 vs. (+) siControl. P<0.05^§^, 0.01^§§^, 0.001^§§§^.* **[B] Cell viability of BEAS-2B cells upon cigarette smoke exposure in both SCAL1-transfected and non-transfected groups by MTT assay.** [a’] Viability of BEAS-2B cells exposed to 20% CSE only. [b’] Viability of BEAS-2B cells exposed to 20% CSE and antisense SCAL1. [c’] Viability of BEAS-2B cells exposed to 20% CSE and scrambled siRNA as siControl. [d’] Comparison of the viability of both siSCAL1 (+) and siSCAL1 (-) BEAS-2B cells with siControl treated BEAS-2B cells exposed to 20% CSE for 2 hours, 4 hours, and 6 hours followed by 20 hours recovery. *Error bars represent SEM. The results depicted represent the mean of 3 biological replicates and technical replicates. P<0.05*, 0.01**, 0.001*** (CSE treated vs. Untreated), P<0.05^#^, 0.01^##^, 0.001^###^ (between-group, non-transfected), P<0.05*^θ^*, 0.01*^θθ^*, 0.001*^θθθ^ *(siSCAL1-4hour vs. siControl-4hour), P<0.05^§^, 0.01^§§^, 0.001^§§§^ (siSCAL1-6hour vs. siControl-6hour), P<0.05^@^, 0.01^@@^, 0.001^@@@^ (between-group; transfected vs. non-transfected)*.

### Increased cell viability on CSE exposure

The siSCAL1 (-) BEAS-2B cells had significantly increased cell proliferation and survival after 6 hours of incubation in 20% CSE compared to the other CSE incubation hours. Contrarily, siSCAL1 (+) cells had reduced cell proliferation and cell viability compared to the siSCAL1 (-) cells for all CSE exposed and unexposed groups. However, siControl (+) had no significant difference from siSCAL1(-) cells (**Fig. 2[B]**).

### Changes in Cellular Morphology on CSE Exposure

In the siSCAL1 (-) CSE (+) group, we observed a heterogeneous population of large, flattened, amoeboid-like multinucleated cells forming cone-shaped migrating fronts, prominent focal adhesions and intercellular connections or stress fibers resembling tunneling nanotubes. The cells lost contact inhibition and had grown in multiple layers, forming clusters of diffusely distributed cells at higher incubation in CSE compared to siSCAL1 (-) CSE (-) cells (**Fig. 3[a-f]**). However, siSCAL1 (+) BEAS-2B cells, upon 20% CSE treatment, appeared to be rounded at 2 hours of CSE incubation, while at 4 hours and 6 hours of CSE incubation, most cells died. The survived cells appeared with epithelial morphology, regular-shaped nuclei, and few focal adhesions and stress fibers (**Fig. 3[a’-d’]**). The siControl set showed no significant change from siSCAL1 (-) BEAS-2B cells.

**Figure 3.**
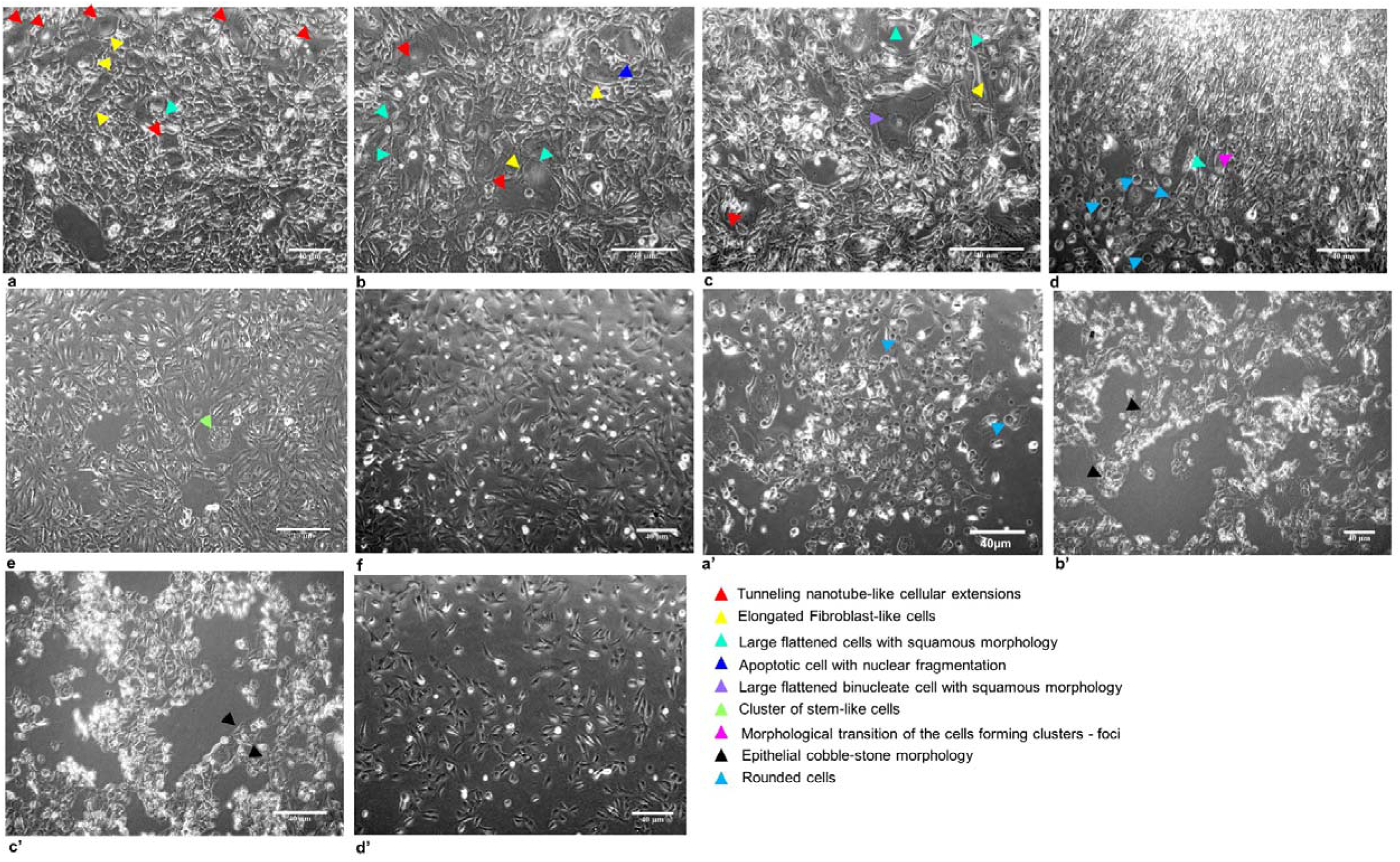
Comparative cellular morphology of BEAS-2B cells. **[a] siSCAL1 (-), CSE (+) for 2 hours, [b] siSCAL1 (-), CSE (+) for 4 hours, [c-e] siSCAL1 (-), CSE (+) for 6 hours, [f] siSCAL1 (-), CSE (-), [a’] siSCAL1 (+), CSE (+) for 2 hours, [b’] siSCAL1 (+), CSE (+) for 4 hours, [c’] siSCAL1 (+), CSE (+) for 6 hours, [d’] siSCAL1 (+), CSE (-)**. *The magnification of images taken is 400X*.

### SCAL1 mediated ROS mitigation and cytoprotection in BEAS-2B cells

We observed no significant difference in the intracellular ROS levels between siSCAL1 (-) CSE (+) cells at 6 hours of incubation and siSCAL1 (-) CSE (-) cells. At 2- and 4 hours of incubation of siSCAL1 (-) cells in CSE, high intracellular ROS was detected compared to siSCAL1 (-) CSE (-) cells. In siSCAL1 (+) CSE (+) BEAS-2B cells, we found higher levels of intracellular ROS compared to siSCAL1 (-) CSE (+) BEAS-2B cells for all CSE exposure time points, as shown in **Fig. 4 [A-D, a-d, a’-d’]**. The siControl (+) cells also exhibited a comparable pattern to siSCAL1 (-) CSE (+) cells.

**Figure 4.**
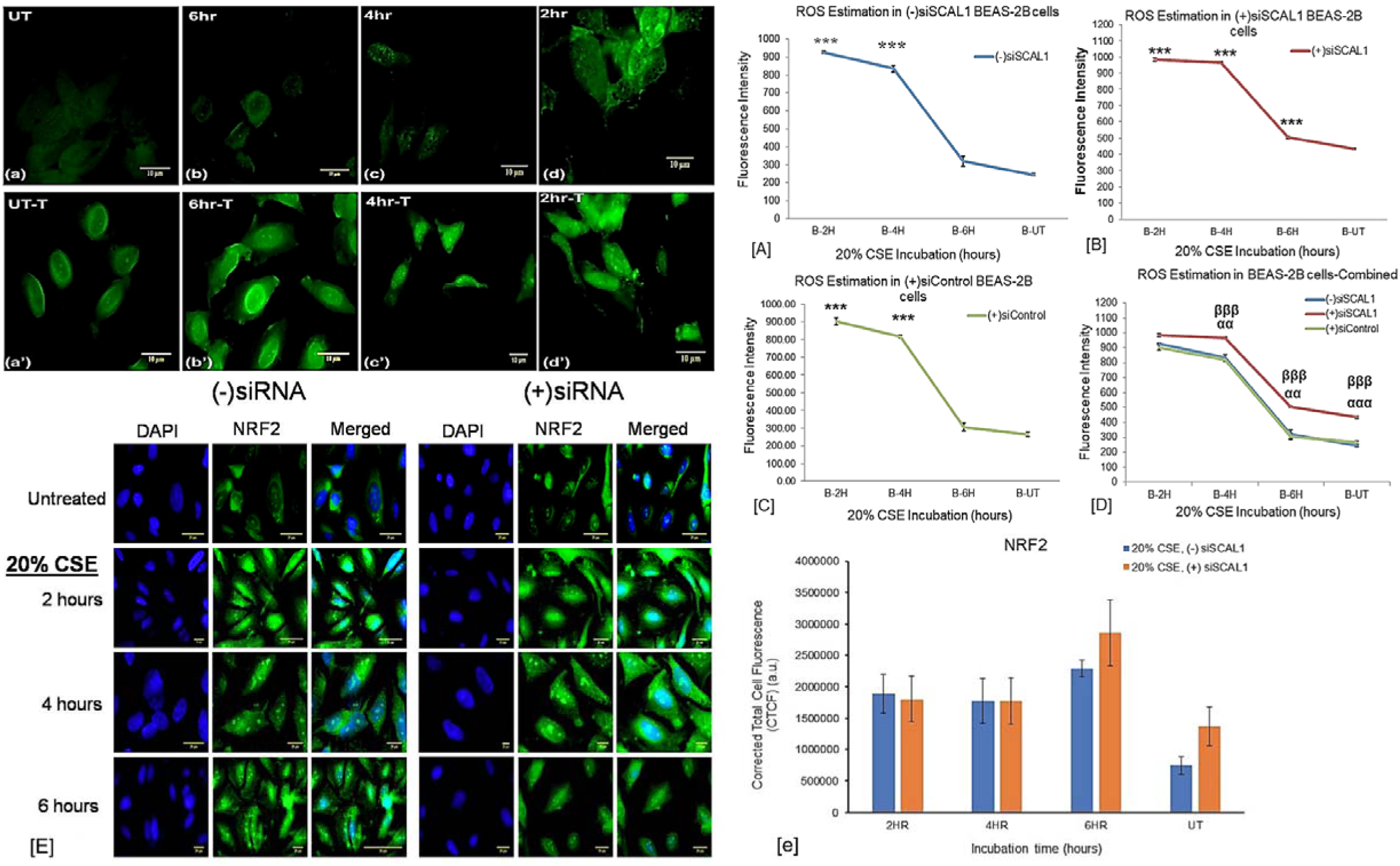
Estimation of 20% CSE-induced ROS in BEAS-2B cells by H2-DCFDA assay. **[A] siSCAL1 (-) 20% CSE (+) for 2, 4, and 6 hours**, **[B] (+) siSCAL1 and 20% CSE (+) for 2, 4, and 6 hours, [C] (+) siControl 20% CSE (+) for 2, 4, and 6 hours, [D] Combined representation**. Fluorescence imaging was also done that qualitatively depicted the ROS levels in; **(a) siSCAL1 (-) and CSE (+) for 2 hours, (b) siSCAL1 (-) and CSE (+) for 4 hours, (c) siSCAL1 (-) and CSE (+) for 6 hours, (d) siSCAL1 (-) and CSE (-)**. Similarly, **(a’) siSCAL1 (+) and CSE (+) for 2 hours, (b’) siSCAL1 (+) and CSE (+) for 4 hours, (c’) siSCAL1 (+) and CSE (+) for 6 hours, (d’) siSCAL1 (+) and CSE (-).** Error bars represent SEM. *p<0.05*, 0.01**, 0.001\*\*\** (comparison between 20% CSE exposed vs 20% CSE unexposed). *p<0.05*^α^*, 0.01*^αα^*, 0.001*^ααα^ (comparison between siSCAL1 treated vs siSCAL1 untreated cells between same hours of 20% CSE incubation), and *p<0.05*^β^*, 0.01*^ββ^*, 0.001*^βββ^ (comparison between siSCAL1 treated vs siControl untreated cells between same hours of 20% CSE incubation). The scale bar is 10μm. **[E] Expression of NRF2 for 20% CSE exposure in both siSCAL1 (-) and siSCAL (+) BEAS-2B cells. [e] The Corrected Total Cell Fluorescence (CTCF) of siSCAL (-) CSE (+) vs. siSCAL (+) CSE (+) cells for 2, 4, and 6 hours of CSE incubation and between siSCAL1 (-) CSE (-) and siSCAL1 (+) CSE (-) cells.** The representative images were produced, and the CTCF was done using the mean of at least three biological replicates. Error bars represent SEM. Scale 20 μm. *p<0.05*, 0.01**, 0.001\*\*\** (comparison between CSE-exposed vs CSE-unexposed groups).

Since NRF2, a key regulator of cytoprotective genes, plays a crucial role in inducing cytoprotective genes through the *NRF2*/antioxidant response element pathway, we have evaluated the expression of NRF2 in our study groups.

In siSCAL1 (-) BEAS-2B cells, we found 1.69-, 1.77-, and 1.99-fold upregulation of NRF2 upon exposure to 20% CSE for 2 hours, 4 hours, and 6 hours, respectively (**Fig. 6**). There was no significant difference in the CTCF of NRF2 between siSCAL1 (-) and siSCAL1 (+) BEAS-2B cells matched with CSE exposure groups. (**Fig. 4 [E-e]**).

### SCAL1-mediated cancer stemness and tumorigenic differentiation in BEAS-2B cells Cancer stemness CD133

We found significant 4.72-, 5.93, and 4.54-fold overexpression of CD133 (133 kD) for 2, 4, and 6 hours of incubation, respectively, in siSCAL1 (-) CSE (+) BEAS-2B cells compared to siSCAL1 (-) CSE (-) BEAS-2B cells. Similarly, we found 1.98-, 2.96, and 3.20-fold overexpression (*p*<0.05) of CD133 (95 kD) for 2, 4, and 6 hours of incubation, respectively (**Fig. 6**). However, siSCAL1 (+) BEAS-2B cells exposed to similar CSE exposures exhibited significant downregulation of CD133 (133 kD) and CD133 (95 kD) at all incubation hours.

We observed significantly higher CTCF of CD133 expression in siSCAL1 (-) CSE (+) BEAS-2B cells compared to the CSE (-) group. There was a significant reduction in the CTCF of CD133 in siSCAL1(+) cells compared to the siSCAL1(-) cells matched with CSE exposure groups (**Fig. 5 [A-a]**).

**Figure 5.**
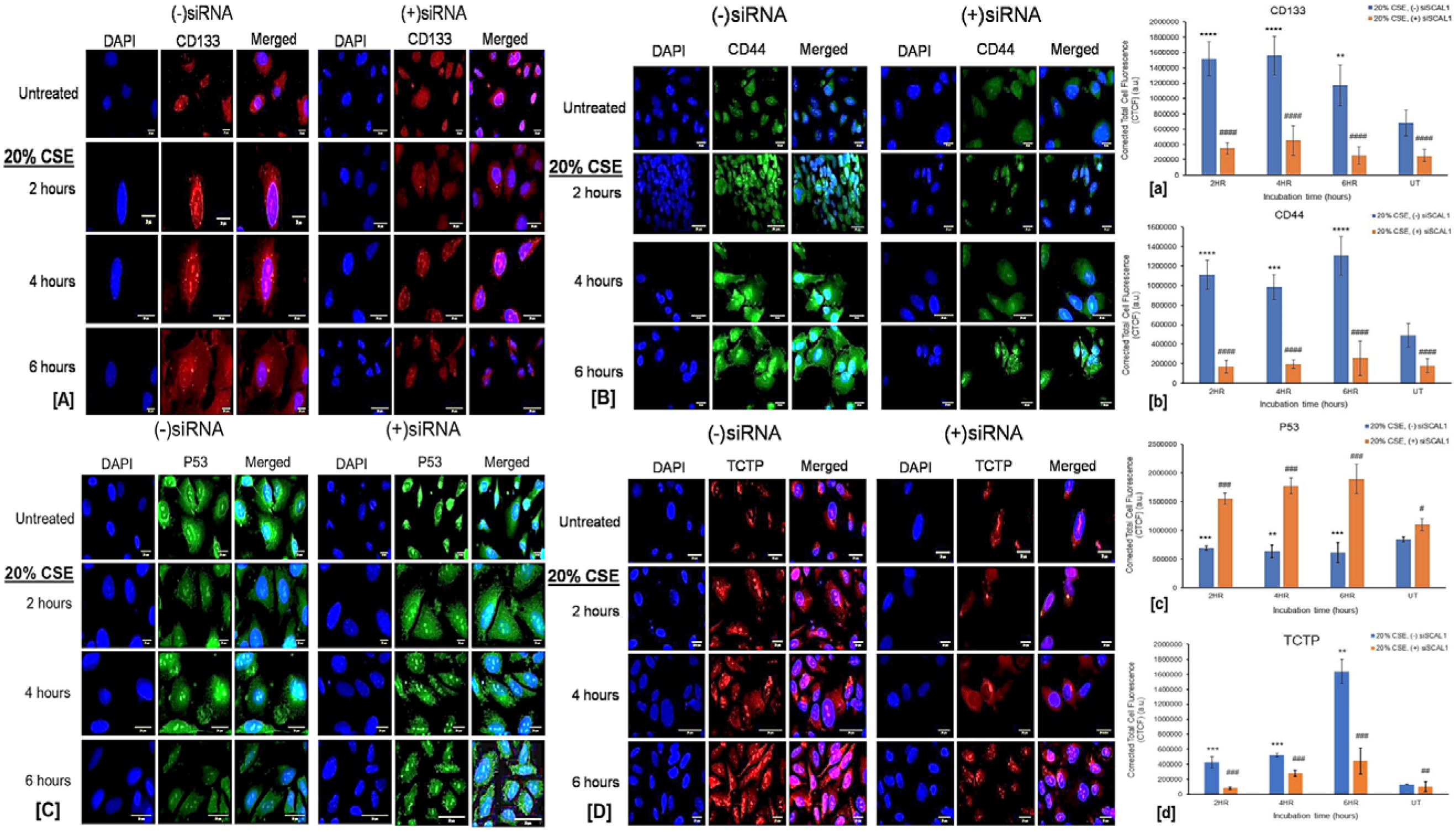
Comparative expression and maintenance of [A] CD133 (CSC marker), [B] CD44, [C] P53, and [D] TCTP in siRNA/siSCAL1 (-) CSE (-) with siSCAL1 (+) CSE (-), siSCAL1 (-) CSE (+) with siSCAL1 (+) CSE (+) for 2, 4, and 6 hours of incubation in CSE. The corresponding CTCF assessments for [a] CD133, [b] CD44, [c] P53, and [d] TCTP are represented in bar graphs. *The representative graphs were produced using the mean of at least three biological replicates. Error bars represent SEM. The scale is 20* μ*m. p<0.05*, 0.01**, 0.001*** (Comparison between CSE-exposed vs CSE-unexposed), P<0.05^#^, 0.01^##^, 0.001^###^ (comparison between siSCAL1 treated vs siSCAL1 untreated cells)*.

### CD44

After exposure to 20% CSE, siSCAL1 (-) BEAS-2B cells exhibited a significant upregulation of 2.48-, 2.39-, and 2.55-folds for the CSC marker CD44 (80 kD) at 2, 4, and 6 hours of incubation, respectively, compared to siSCAL1 (-) CSE (-) BEAS-2B cells (**Fig. 6**). Conversely, siSCAL1 (+) cells showed a significant downregulation matched with the respective CSE exposure groups.

**Figure 6.**
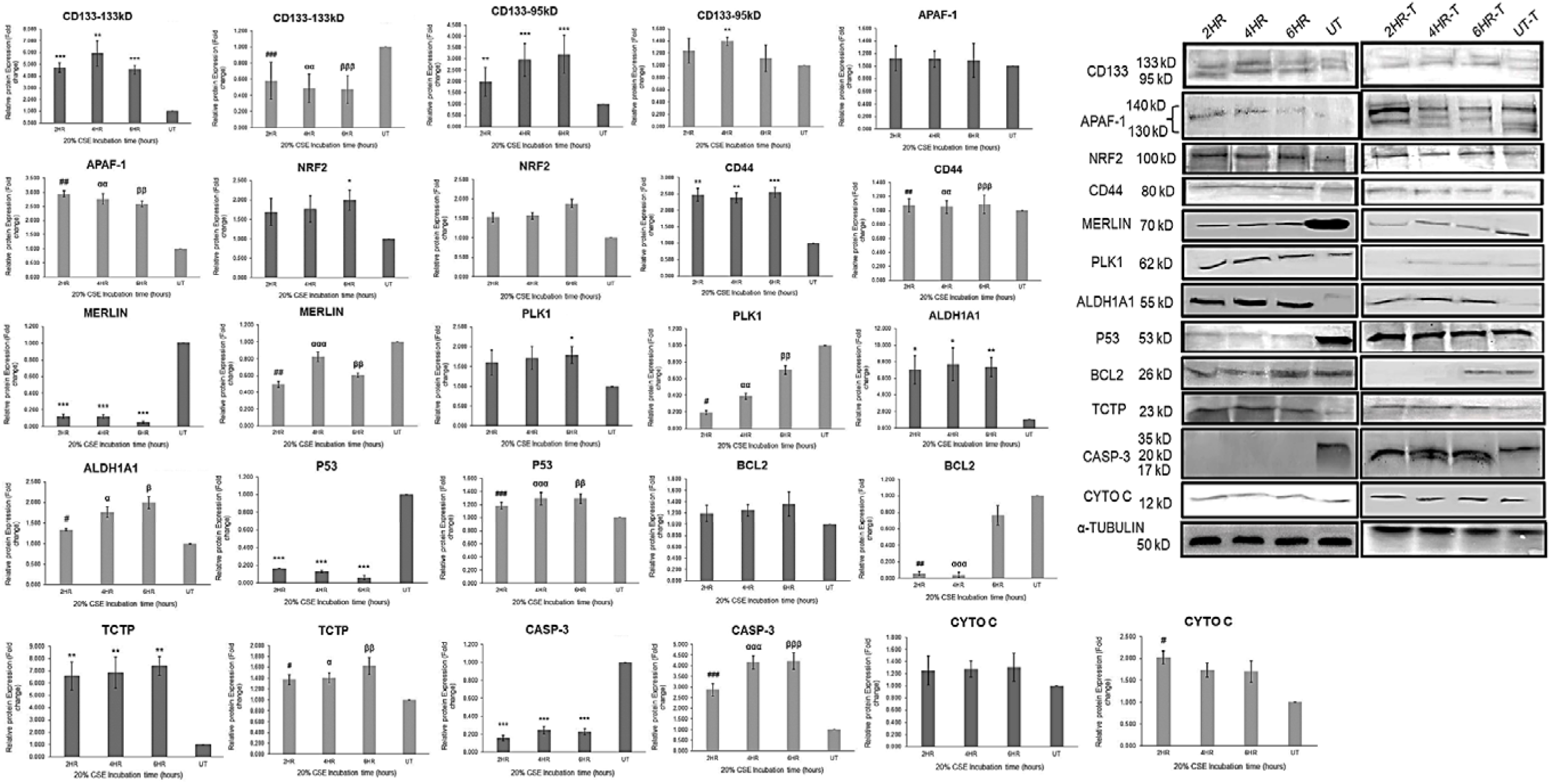
Relative expression of marker proteins normalized to α-tubulin expression for cytoprotection, cancer stemness, apoptosis, cell proliferation, and cytoskeletal organization responsible for tumorigenic differentiation of BEAS-2B cells. The comparisons were done between **siSCAL1 (-) CSE (-) with siSCAL1 (+) CSE (-), siSCAL1 (-) CSE (+) with siSCAL1 (+) CSE (+) for 2, 4, and 6 hours of incubation in CSE.** The densitometry was done using the mean of at least three biological replicates. Error bars represent SEM. *p<0.05*, 0.01**, 0.001\*\*\** (Comparison between CSE-exposed vs CSE-unexposed), *p<0.05*^β^*, 0.01*^ββ^*, 0.001*^βββ^ (comparison between siSCAL1 treated vs siSCAL1 untreated cells between 4 hours of 20% CSE incubation), *p<0.05*^α^*, 0.01*^αα^*, 0.001*^ααα^ (comparison between siSCAL1 treated vs siSCAL1 untreated cells between 2 hours of 20% CSE incubation), and *p<0.05^#^, 0.01^##^, 0.001^###^* (comparison between siSCAL1 treated vs siSCAL1 untreated cells between 2 hours of 20% CSE incubation).

We observed significantly higher CTCF of CD44 (80 kD) in siSCAL (-) CSE (+) BEAS-2B cells compared to the siSCAL (-) CSE (-) cells. We found significantly lower CTCF of CD44 in siSCAL1(+) cells matched with CSE exposures (**Fig. 5 [B-b]**).

### ALDH1A1

Our data revealed a 7.033-, 7.704-, and 7.365-fold overexpression of ALDH1A1 (55 kD) in 2, 4, and 6 hours of CSE incubation, respectively, in siSCAL1(-) cells compared to siSCAL1 (-) CSE (-) cells. A significant decline in ALDH1A1 expression was observed in siSCAL1 (+) cells matched to respective CSE exposures (**Fig. 6**).

## Tumorigenic differentiation

### P53-TCTP reciprocal expression

We found a significant decrease (p<0.05) in P53 and a significant increase (p<0.05) in the translationally controlled tumor protein (TCTP, 23 kD) in siSCAL1 (-) CSE (+) BEAS-2B cells for all incubation time points compared to siSCAL1 (-) CSE (-) cells. However, in siSCAL1 (+) cells, we found a significant upregulation of P53 and a significant downregulation of TCTP matched for the respective CSE exposures (**Fig. 6**). We observed similar trends in the outcomes through immunofluorescence imaging for the groups mentioned above (**Fig. 5 [C-c]**).

### PLK1

Our findings showed a 1.60-, 1.72-, and 1.79-fold overexpression of PLK1 in siSCAL1 (-) CSE (+) BEAS-2B cells exposed for 2, 4, and 6 hours, respectively, compared to CSE (-) cells. Significant upregulation was observed in 6 hours of CSE incubation compared to CSE (-) cells. However, a significant decline (p<0.05) in the expression of PLK1 was seen in siSCAL1 (+) BEAS-2B cells matched for similar CSE exposures (**Fig. 6**).

### MERLIN (NF2)

We found a statistically significant downregulation of Merlin of 0.12-, 0.118-, and 0.051-fold in siSCAL1 (-) CSE (+) BEAS-2B cells for 2, 4, and 6 hours of exposures, respectively, compared to the siSCAL1 (-) CSE (-) cells. Interestingly, siSCAL1 (+) cells matched with similar CSE exposures showed a significant increase in the expression of Merlin compared to the siSCAL1 (-) cells (**Fig. 6**).

### Spheroid Formation-Cancer stemness and tumorigenicity of BEAS-2B cells

We found significantly (*p*<0.05) increased number of spheroid colonies on the 11th and 21^st^ day from baseline in the siSCAL1 (-) CSE (+) BEAS-2B cells for 4 and 6 hours compared to CSE (-) cells (**Fig. 7 [A-C, A’-C’, D-E]**). However, the number of spheroids significantly reduced in siSCAL1 (+) BEAS-2B cells matched for similar CSE exposures in the siSCAL1 (-) cells (**Fig. 7 [A’’-C’’, A’’’-C’’’, D-E]**). The trend of spheroid formation in siSCAL1 (+) and siSCAL1 (-) cells shows concordance with the expression of CSC markers, like CD133, CD44, and ALDH1A1 in these two groups of BEAS-2B cells matched for similar CSE exposures.

**Figure 7.**
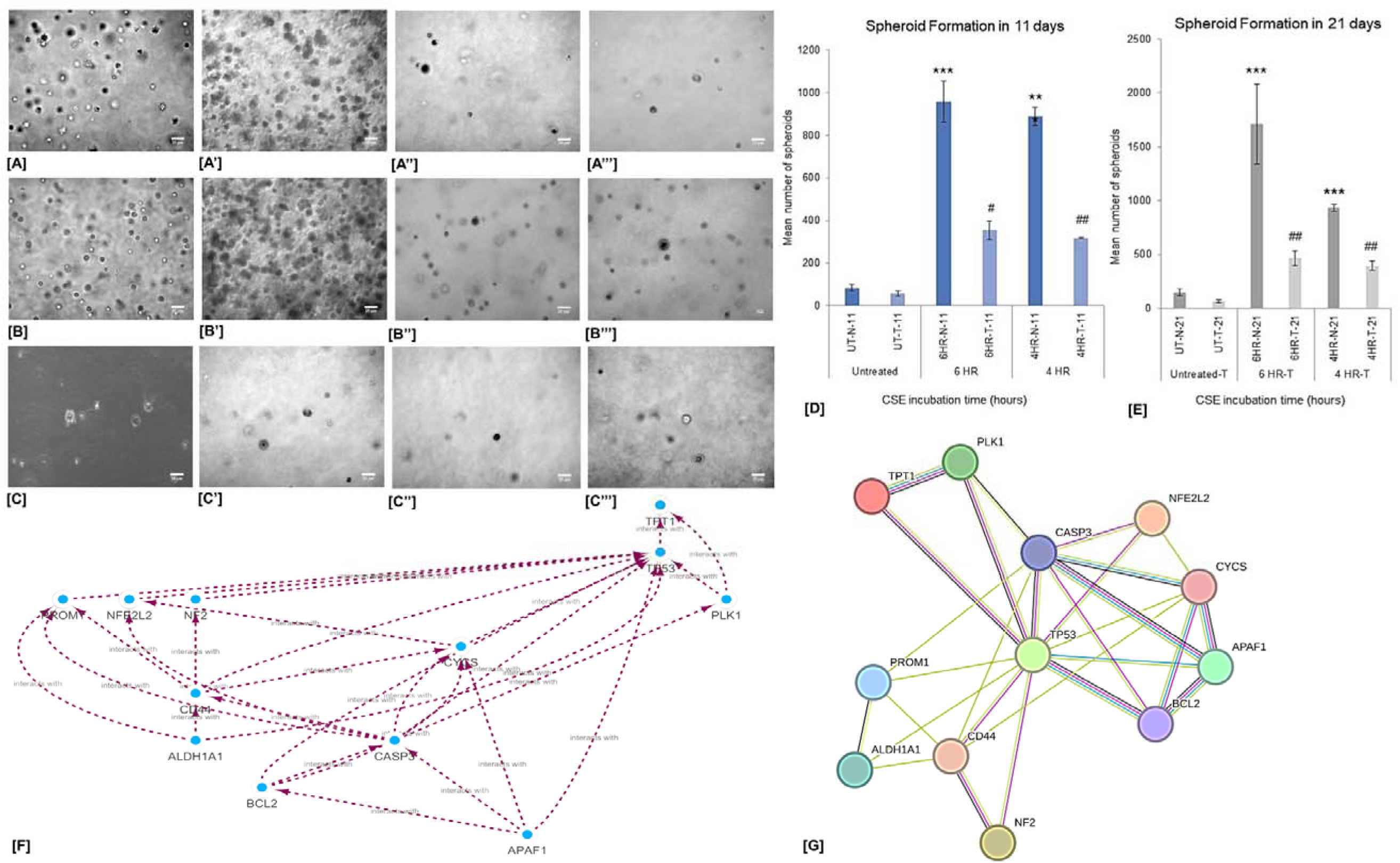
**[A] Number of spheroids at 4 hours of CSE incubation at 11 days from cell seeding in siSCAL1 (-) BEAS-2B cells, [A’] Number of spheroids at 4 hours of CSE incubation at 21 days from cell seeding in siSCAL1 (-) BEAS-2B cells, [A’’] Number of spheroids at 4 hours of CSE incubation at 11 days from cell seeding in siSCAL1 (+) BEAS-2B cells, [A’’’] Number of spheroids at 4 hours of CSE incubation at 21 days from cell seeding in siSCAL1 (+) BEAS-2B cells. [B] Number of spheroids at 6 hours of CSE incubation at 11 days from cell seeding in siSCAL1 (-) BEAS-2B cells, [B’] Number of spheroids at 6 hours of CSE incubation at 21 days from cell seeding in siSCAL1 (-) BEAS-2B cells, [B’’] Number of spheroids at 6 hours of CSE incubation at 11 days from cell seeding in siSCAL1 (+) BEAS-2B cells, [B’’’] Number of spheroids at 4 hours of CSE incubation at 21 days from cell seeding in siSCAL1 (+) BEAS-2B cells. [C] Number of spheroids for CSE (-) at 11 days from cell seeding in siSCAL1 (-) BEAS-2B cells, [C’] Number of spheroids for CSE (-) at 21 days from cell seeding in siSCAL1 (-) BEAS-2B cells, [C’’] Number of spheroids for CSE (-) at 11 days from cell seeding in siSCAL1 (+) BEAS-2B cells, [C’’’] Number of spheroids for CSE (-) at 21 days from cell seeding in siSCAL1 (+) BEAS-2B cells. [D-E] Mean number of spheroids at 11 and 21 days from cell seeding for siSCAL1 (-) and siSCAL1 (+) BEAS-2B cells.** Error bars represent SEM. *p<0.05*, 0.01**, 0.001\*\*\** (Comparison between CSE-exposed vs CSE-unexposed), *p<0.05^#^, 0.01^##^, 0.001^###^* (comparison between siSCAL1 treated vs siSCAL1 untreated cells between 2 hours of 20% CSE incubation). **[F-G] Protein-Protein Interactome of the markers from STRING v10.0**

Interestingly, after 21 days, we observed an increase in the invasiveness or spread of the siSCAL1 (-) cells compared to the siSCAL1 (+) cells matched for similar CSE exposures. The round cells changed their architecture to elongated spindle-shaped structures spreading out from the spheroids, forming tunneling nanotube-like structures.

### Apoptosis

We assessed the expression of key apoptotic markers in siSCAL1(-) and siSCAL1 (+) BEAS-2B cells under 20% CSE exposure for 2, 4, and 6 hours of incubation.

### APAF-1

We observed no significant difference in the expression of APAF-1 between siSCAL1 (-) CSE (+) and siSCAL1 (-) CSE (-) cells. APAF-1 was significantly overexpressed by 2.94-, 2.76-, and 2.59-fold (p<0.05) in siSCAL1 (+) BEAS-2B cells with CSE (+) for 2, 4, and 6 hours of incubation, respectively, compared to the siSCAL1(-) cells matched for similar CSE exposures (**Fig. 6**).

### CYTOC

We found no significant difference in the expression of CYTOC (12 kD) between siSCAL1 (-) CSE (+) and siSCAL1 (-) CSE (-) BEAS-2B cells. We found a significantly higher expression of CYTOC in siSCAL1 (+) cells at 2 hours of CSE exposure compared to siSCAL1 (-) cells for the same CSE exposure. For 4- and 6-hours of CSE exposure, siSCAL1 (+) cells showed higher expression of CYTOC compared to siSCAL1 (-) BEAS-2B cells (**Fig. 6**).

### BCL2

We found a 1.19-, 1.25-, and 1.36-fold expression of the anti-apoptotic BCL2 in siSCAL1 (-) BEAS-2B cells with CSE (+) for 2, 4, and 6 hours, respectively, compared to the CSE (-) cells. In siSCAL1(+) CSE (+) BEAS-2B cells, we found a significant downregulation of BCL2 at 6 hours compared to CSE (-) cells. However, we did not find any noticeable expression of BCL2 at 2 and 4 hours in siSCAL1 (+) cells (**Fig. 6**).

### CASP-3

Our data showed expression of uncleaved caspase-3 (35 kD) in the siSCAL1 (-) CSE (-) BEAS-2B cells, while no expression of caspase-3 was found at 2, 4, and 6 hours of incubation in siSCAL1 (-) CSE (+) cells. Interestingly, in siSCAL1 (+) BEAS-2B cells, we found the expression of cleaved caspase-3 (20 kD and 17 kD) at 2, 4, and 6 hours of CSE incubation (**Fig. 6**).

### Protein-Protein Interaction Analysis

We observed TP53 as the central player in the interactome with experimental evidence of protein-protein interaction with TCTP, PLK1, CASP3, NFE2L2, MERLIN (NF2), BCL2, and CD44. STRING showed experimental evidence of protein-protein interaction between TCTP—PLK1, CD44—MERLIN, CASP3—APAF1, CASP3—NFE2L2, and CASP3—BCL2 (**Fig. 7[F-G]**). Moreover, enrichment analysis identified the top enriched pathways, processes, genes, compartments, and tissues associated with apoptosis, cell death, and cellular stress response. This emphasizes the statistically significant enrichments, with lower p-values indicating stronger associations. The results indicate apoptosis induction via p53 signaling and cellular stress in cancer/blood cell lines. The key genes TP53, CYCS, CASP3, BCL2, and APAF1 are involved in apoptosis, p53 signaling, and intrinsic apoptotic pathways. This highlights how these genes connect to the central enriched processes related to apoptosis activation and cell death.

## Discussion

The study provides novel insights into the function of the long non-coding RNA SCAL1 in BEAS-2B cells when exposed to 20% CSE and its potential impact on lung cancer development. The study investigates the effects of SCAL1 on cell viability, cytoprotection, cancer stemness, tumorigenic differentiation, cellular invasiveness, and apoptosis regulation.

Tobacco smoking leads to oxidative damage and the acquisition of genetic lesions, with the formation of DNA adducts [21–23]. However, the mechanisms by which cells survive and remain viable despite these incessant genetic insults remain unclear. More recently, lncRNAs were proposed to play a role in detoxification processes as a response to xenobiotic insults. The metabolism of tobacco smoke plays a vital role in smoke-induced toxicity, oxidative stress, and subsequent DNA damage [24]. The smoke metabolites can impart various damaging effects on the lung cells that could lead to cellular transformations and the inception of tumorigenic differentiation. Phase I metabolism genes function in the bio-activation of smoke components, such as aryl hydrocarbon receptor (*AhR*) [25–27] and Cytochrome P450 2A6 (*CYP2A6*) [28, 29], into potent carcinogenic metabolites, imparting positive carcinogenic stress to the tobacco smoke-exposed lung [30].

Thai et al. [10] found that SCAL1 levels were considerably higher in airway epithelia of smokers compared to non-smokers upon analyzing global RNA expression in smoke-exposed lung cancer cells. In one dataset, the levels were up to 5.3 times higher, while in another, they were up to 3.9 times higher. In our study, SCAL1 upregulation in BEAS-2B cells on 20% CSE exposure for 6 hours shows collinearity to its expression in metastatic lung cancer cell line CL1-5 (i.e., 15-fold compared to CL1-0 non-metastatic lung cancer cell line) while the expression of SCAL1 at 2 hours shows similarity to that of the bronchial epithelium of smokers. Thus, it can be concluded that exposure to 20% CSE stimulates SCAL1 expression in BEAS-2B cells. The silencing of SCAL1 would reveal its biological role in lung carcinogenesis and progression. A parallel experiment with siSCAL1 (+) cells with 20% CSE exposure was conducted to establish its role.

In this study, the results suggest an essential role of the lncRNA SCAL1 in maintaining cell viability and survival under CSE-induced oxidative stress. It seems that BEAS-2B cells adapt to oxidative stress, undergo cellular transformation, and show higher survival under prolonged exposure to cigarette smoke extract. Higher cell survival indicates cytoprotection and enhanced cell proliferation due to SCAL1 upregulation since SCAL1 was previously reported to be cytoprotective [10]. Our data also showed increased cell survival and viability under CSE-induced SCAL1 upregulation. Another report stated that chronic exposure of BEAS-2B cells with CSE induces tumorigenic transformation with hyperplastic growth, loss of contact inhibition, aberrant and condensed nuclei, and altered nucleocytoplasmic ratio [31]. These findings indicate an adaptation of the BEAS-2B cells to the CSE, followed by their hyperplastic growth on prolonged exposure to 20% CSE, like the earlier report [31]. Our data suggest a role of SCAL1 in the regulation of cytoskeletal dynamics of the cell that altered the native epithelial morphology of BEAS-2B cells and formed populations of fusiform, enlarged, flattened, and round cells under oxidative and hypoxic stress generated by CSE exposure. The resultant morphology of the CSE-exposed BEAS-2B cells indicates tumorigenic transformation, which has been reported earlier [31]. siSCAL1 (-) BEAS-2B cells treated with 20% CSE resulted in both common and distinct temporary structures related to cell migration and invasive capability. They showed the formation of filamentous extensions resembling tunneling nanotubes, as Cruz et al. [32] reported. These intercellular connections usually allow the passage of organelles, plasma membrane components, and cytoplasmic molecules between connected cells and may further promote invasiveness [33–35]. These filamentous extensions play a role in cell-to-cell communication and transfer of cellular cargo that is believed to be involved in tumor invasion and metastasis [36]. Such intercellular extensions create specific communication pathways and establish open-ended channels that maintain membrane continuity to help balance stress factors caused by pathological changes and changing conditions such as oxidative stress or nutrient shortage. The intercellular transfer via filamentous extensions leads to the emergence of molecular pathways that significantly affect critical physiological processes like cell survival, redox, or metabolic homeostasis via ROS detoxification and mitochondrial heteroplasmy [37, 38]. The development of filamentous extensions in CSE-exposed cells may be a way to adapt to oxidative stress caused by smoking, is commonly seen as a positive indicator for lamellipodia formation, and may play a part in the nucleus’ malleability and deformability during cellular migration [39].

An earlier study showed high expression of embryonic stem cell markers in airway epithelial cells of smokers [3], which led to the idea of the role of tobacco smoke in the dysregulation of the cell fate and differentiation pathways contributing to lung carcinogenesis. Exposure of non-tumorigenic cells to high ROS generally increases the expression of P53 and decreases the NRF2 levels, leading to oxidative stress-induced apoptosis and decreasing cell survival [40]. Cigarette smoking and lung carcinogenesis are intricately linked, and the accumulation of smoke metabolites is thought to generate cells with stem properties, called cancer stem cells. SCAL1 was overexpressed in aggressive lung cancer cell lines and healthy bronchial epithelial cells in response to cigarette smoke exposure. It is transcriptionally regulated by the nuclear factor erythroid 2-related factor (NRF2) kelch-like ECH-associated protein 1 (KEAP1). The knockdown of SCAL1 potentiated cellular cytotoxicity induced by CSE *in vitro* [10].

The study established the role of SCAL1 as an important biomarker for detecting tumorigenic differentiation and the induction of cancer stem cell properties in healthy human bronchial epithelial cells in response to cigarette smoke exposure. NRF2 is a potent transactivator and is upregulated in response to oxidative stress induced by cigarette smoke. It controls the expression and activity of several genes involved in redox homeostasis and induces pro-survival signals [41]. On the contrary, TP53 behaves as the guardian of the genome, with its bifunctional pro-oxidant and antioxidant properties. Low levels of reactive oxygen species (ROS) induce p21-dependent upregulation of antioxidant genes like *NRF2* and its target genes, resulting in the mitigation of ROS/electrophile-induced damages in the cells. [42], mainly belonging to the lung epithelium. Exposure of cells to a high amount of ROS results in enhanced p53 expression, causing p53-dependent apoptosis. This indicates a bimodal regulation between p53 and NRF2 that could influence the tumor suppressive function of p53 by coordinated regulation between survival and apoptotic pathways [43]. The data showed upregulation of NRF2 in BEAS-2B cells upon 20% CSE exposure. However, siRNA-mediated silencing of SCAL1 was not found to affect the expression of NRF2. Thus, it indicates the lack of any feedback regulation of NRF2 by SCAL1.

Interestingly, negative feedback signaling between p53 and TCTP (*Translationally controlled tumor protein*) [44] has been found to regulate downstream Akt pathway signal transduction that enhances the metabolism level, proliferation, metastasis, and survival of lung cancer cells from oxidative stress-induced apoptosis [45]. Recent findings suggest that overexpression of TCTP attenuates p53 expression and further enhances its proteasomal degradation through *Mdm2*. TCTP has also been linked with phenotypic reprogramming and modification of cancer stem cell compartments in breast cancer [46]. The data above corroborates with the study’s finding that upregulation of SCAL1 mediates the downregulation of p53 and upregulation of TCTP in BEAS-2B cells, suggesting a probable tumorigenic differentiation in the otherwise non-tumorous BEAS-2B cells. We observed the recovery of p53 expression and a decrease in TCTP expression in siSCAL1 (+) BEAS-2B cells. Interestingly, the similar downregulation of TCTP in cancer cells has been correlated with the decline in cell proliferation and apoptotic targeting [44]. The downregulation of TCTP has been associated with increased chemosensitivity of HeLa cells in response to etoposide treatment [47].

The induction of the expression of cancer stem cell markers, such as CD133 and CD44, is required to develop tumorigenicity in the normal bronchial epithelial cells. CD133 is a well-characterized tumor-initiating CSC marker in various cancers, including lung cancer. The *CD133* promoter P1 was reported to have a non-canonical p53-binding sequence where p53 directly binds and recruits Histone Deacetylase 1 (HDAC1), leading to negative regulation of *CD133* expression [48]. This suggests the impact of p53 in regulating the growth and tumorigenic potential of cancer stem cells by CD133 suppression [48]. Glycosylation of CD133 (Prominin-1) is reported to be essential in stem cell maintenance [49]. De-glycosylation was observed with the differentiation of stem cells without affecting its mRNA and protein expression [50]. The data indicates similar p53-dependent regulation of *CD133* expression in 20% CSE-exposed non-tumorigenic BEAS-2B cells. The siRNA-mediated silencing of SCAL1 was found to upregulate P53, and subsequent downregulation of CD133 was evident from the data mentioned in the result section. Thus, it can be inferred that SCAL1 mediates the p53-dependent CD133 expressional regulation.

Similarly, the CD44 promoter also harbors non-canonical p53-binding sites where p53 directly binds and contributes to the repression of CD44 expression, leading to the inhibition of tumor cell survival [51]. Switching of the expression of CD44v isoform in epithelial cells to CD44s (standard isoforms; M.W.∼80kD) in mesenchymal cells is the central mediator of invadopodium formation during epithelial-to-mesenchymal transition (EMT) in breast cancer invasion and metastasis [52]. CD44 has been reported to actively participate in the maintenance of stem cell population niches by inducing dormancy and resistance to apoptosis [53]. CD44 is a potential marker of CSCs in lung cancer and plays a role in various cellular processes, including metastasis, EMT, and immune cell infiltration [54–56]. Like CD133, SCAL1 is a critical player in the p53-mediated regulation of CD44 expression in 20% CSE-exposed non-tumorigenic BEAS-2B cells. The co-expression of CD44 and CD133 has been observed in non-small cell lung cancer cells. These cells exhibit DNA damage response pathways and dormant polyploid giant cancer cell enrichment, possibly related to their p53 status [57].

According to a recent meta-analysis, increased expression of ALDH1A1 is linked to poor overall survival (OS) and disease-free survival in lung cancer patients [58]. Patients with positive ALDH1A1 expression were ∼ twice more likely to suffer from relapse than those with negative ALDH1A1 expression [59]. In addition, the expression of ALDH1A1 was positively correlated with the stage and grade of lung tumors and related to a poor prognosis for patients with early-stage lung cancer [59]. Interestingly, our study showed overexpression in siSCAL (-) BEAS-2B cells treated with 20% CSE compared to siSCAL (+) cells, indicating a role of SCAL1 in regulating ALDH1A1 expression, which is again an important cancer stem cell marker.

Our study revealed that siSCAL1 (-) cells exposed to 20% aqueous CSE for 4 and 6 hours promoted spheroid formation, while siSCAL1 (+) BEAS-2B cells showed a reduction in spheroid formation under similar CSE exposure. The findings also showed a correlation between spheroid formation and the expression of cancer stem cell markers. Additionally, siSCAL1 (-) cells exhibited increased invasiveness with tunneling nanotube-like networks after 21 days, while siSCAL1 (+) cells did not show such morphological transformations or invasiveness. Thus, the study emphasized the critical regulatory role of SCAL1 in spheroid formation and cellular architecture and behavior in response to CSE exposure.

PLK1 (Polo-like kinase 1) is a crucial regulator of the cell cycle and a vital oncogene in cancer initiation, progression, and drug resistance [60]. It is overexpressed in various human cancers such as glioma, thyroid carcinoma, head and neck squamous cell carcinoma, melanoma, colorectal, oesophageal, ovarian, breast, prostate, and NSCLC [60, 61]. Our study found significant overexpression of PLK1 in siSCAL (-) BEAS-2B cells treated with 20% CSE compared to siSCAL (+) cells, indicating the transformative role of SCAL1 in lung carcinogenesis. Therefore, the overexpression of PLK1 could mediate the increased cell growth, invasiveness, and oncogenic transformation of BEAS-2B cells upon 20% CSE exposure. Inhibition of PLK1 is an attractive strategy to suppress tumorigenic growth and lead to a potential therapeutic approach against lung cancer.

Merlin, encoded by the NF2 (Neurofibromatosis type 2) gene, is a tumor suppressor protein that plays a crucial role in tumorigenesis and metastasis [62–64]. Merlin serves as a linker between transmembrane proteins and the actin cytoskeleton, thereby regulating cytoskeletal dynamics and cell movement [65]. Merlin’s ability to induce contact-dependent growth inhibition suggests its role in regulating cell survival [64]. The loss or inactivation of Merlin has been implicated in various types of human cancers, including lung cancer [66]. Downregulation of the tumor suppressor Merlin in siSCAL1(-) cells that justifies the increased cell proliferation [64] and the alterations in cell shape and structure [67], as observed in the present study. An increase in the expression of Merlin and P53 in siSCAL1-treated cells with similar CSE exposures indicates an increase in apoptosis [68]. Therefore, SCAL1 plays a pivotal role as a potential regulator of multiple hallmarks of lung cancer.

All the proapoptotic proteins showed low expression in siSCAL1 (-) BEAS-2B cells treated with 20% CSE compared to siSCAL1(+) cells, indicating the anti-apoptotic role of SCAL1 in lung cancer. TCTP protects cancer cells from etoposide-induced cell death by inhibiting mitochondria-mediated apoptotic pathways. The interaction of TCTP with Apaf-1 in the Apoptosome engages in the molecular mechanism of TCTP-induced chemoresistance [47]. In our study, the upregulation of TCTP was correlated with the low signal of APAF-1 in immunoblots, which could be due to the interaction of TCTP with APAF-1, generating anti-apoptotic and pro-survival signals in siSCAL1 (-) BEAS-2B cells treated with 20% CSE compared to siSCAL1(+) cells. SCAL1 contributes to cisplatin resistance in human NSCLC by modulating apoptosis, autophagy, migration, and invasion of the cells [69].

The study primarily utilizes the BEAS-2B cell line, an immortalized human bronchial epithelial cell line. While this cell line is commonly used as a model for respiratory research, it may not fully represent the complexity of lung tissue in vivo. Therefore, extrapolating findings from cell culture experiments to the entire lung environment should be done cautiously. Although the study attempts to mimic the effects of cigarette smoke exposure using CSE, it may not fully recapitulate the complexity of *in vivo* exposure to cigarette smoke. CSE lacks the dynamic and diverse mixture of compounds present in actual cigarette smoke, which could influence cellular responses differently. While the study provides valuable insights into the role of the long non-coding RNA SCAL1 in various cellular processes, such as cell viability, cytoprotection, cancer stemness, and apoptosis regulation, the underlying molecular mechanisms remain incompletely understood. Further mechanistic studies are needed to elucidate the precise signaling pathways and molecular interactions involved. Lastly, the study primarily focuses on cellular and molecular changes observed *in vitro*, which may not fully translate to clinical outcomes in human lung cancer patients. Therefore, additional studies, such as clinical trials or epidemiological investigations, are necessary to validate the clinical relevance of the findings.

In this study, we showed the role of SCAL1 in BEAS-2B cells upon 20% aqueous CSE exposure. Our findings reveal that SCAL1 overexpression mitigates ROS-induced oxidative stress, confers cytoprotection, and enhances cancer stem cell-like properties, contributing to cell proliferation and tumorigenic differentiation. Silencing of SCAL1 reverses these effects, underscoring SCAL1’s involvement in lung carcinogenesis and resistance to apoptosis. These insights highlight SCAL1 as a critical regulator in smoke-induced lung carcinogenesis and propose it as a potential therapeutic target for early intervention and treatment of lung cancer linked to smoking.

## Methods

### Cell Cultures

The immortalized human bronchial epithelial (BEAS-2B) cells were cultured in 1:1 v/v mixture of F12 Ham’s and DMEM sterile-filtered (through 0.22μ membrane; PALL) media containing 1% v/v Penicillin/Streptomycin solution (10,000U/ mL; Gibco^TM^), 01% v/v Amphotericin B (Gibco^TM^), 0.06% v/v Gentamycin sulfate (SRL) from 10% w/v stock solution. The BEAS-2B cells are SV40-immortalised human bronchial epithelial cells retaining the attributes of primary cells and previously reported as a suitable model for *in vitro* lung cancer risk assessment that has delayed senescence [70–72]. Complete media was prepared by supplementing with the following growth factors: 1.5% v/v Bovine Pituitary Extract (ThermoFisher Scientific) from 1mg/mL stock solution, 0.02% v/v Recombinant Epidermal Growth Factor (EGF; Abcam) from 10μg/mL stock solution, 0.02% v/v human Transferrin (Merck) from 50mg/mL stock solution, 0.02% v/v albumin solution sterile-filtered (SRL), 0.25% v/v recombinant human Insulin (Abcam) from 2mg/mL stock solution, 0.01% v/v Epinephrine (Abcam) from 5mg/mL stock solution, 0.01% v/v Hydrocortisone (Abcam) from 5mg/mL stock solution, 0.001% v/v Retinoic acid and Triiodo-L-thyronine (T3) (Abcam) each from 1mg/mL stock solutions. The starting cell density of 0.75 x 10^6^- 1 x 10^6^/mL was seeded in 25cm^2^ flasks (Corning®; canted neck and vented cap). The culture was maintained at 37°C in humidified air at 5% CO_2_ and subcultured every 4-5 days before they reached confluency, and the media was changed every second day of cell seeding.

### Preparation of Cigarette Smoke Extract (CSE)

Cigarette smoke extract was prepared using the continuous smoking protocol [73] and the method described by Kode et al. 2006 [74] with slight modifications. The mainstream smoke was continuously sucked into the serum-free complete media through a standard glass tube, and the media was intermittently shaken vigorously 4-5 times for 30secs. A regulated aspiration pump maintained a constant flow rate of 1.050 l/min of cigarette smoke. Two research cigarettes (Kentucky Tobacco R&D Center, Lexington, KY: 3R4F) were used per 10 ml of serum-free complete media as earlier described [10, 75]. The above-obtained media was filter-sterilized through a 0.45μm filter membrane, denoted as 100% CSE. The CSE obtained was used immediately after preparation in proper dilutions (**Fig. 1**).

### Exposure of cultured BEAS-2B cells to CSE

As reported by Thai et al., about 1 x 106 cells/ml of BEAS-2B cells were cultured in complete media until they reached 80-85% confluency and were exposed to CSE. [10] with slight modifications. Cultured BEAS-2B cells were exposed to 20% CSE for 2 hours, 4 hours, and 6 hours. After each incubation hour, the CSE-containing media was aspirated carefully, and new media (complete) was added. The cells were then kept at 37^°^C in a humidified chamber with 5% CO_2_ for 16-20 hours for recovery (**Fig. 1**).

### siRNA Transfection

The SCAL1 siRNA sequences, siSCAL1-1 (CCCACAAAUAGGAAGAAAAdtdt) and siSCAL1-2 (CAUUUCAGUCACUAAAUAAdtdt) used for the study were adopted from the earlier described research [10]. The negative control scrambled siRNA was obtained as a gift from Dr. Somsubhra Nath, Scientist, Basic Research, Saroj Gupta Cancer Centre & Research conducted using Lipofectamine^TM^ RNAiMAX (ThermoFisher scientific; Cat # 13778150) following the modified manufacturer’s protocol (described below) after CSE treatment. The transfection media was removed after 4-6 hours, and complete fresh media was added.

### Cell Viability Assay

BEAS-2B cells were treated with 20% CSE for 2 hours, 4 hours, and 6 hours, followed by media change and 24 hours of recovery. Cell viability was assessed for both siSCAL1(+) and siSCAL1(-) conditions by the CellTiter96® Non-Radioactive Cell Proliferation Assay (MTT assay; Promega, #G4000) following the manufacturer’s protocol.

The experiment was repeated with three sets of cells of the same passage number (biological replicates) and three individual measurements of each biological replicate (technical replicates) for all the treatment conditions. The percentage of cell viability for each treatment type and hour was estimated from the absorbance values. The formula used for the estimation of the rate of cell viability was as follows:

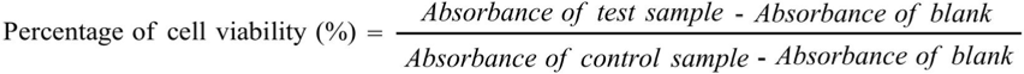

### Microscopy

We evaluated the observable morphological changes in BEAS-2B cells exposed to 20% CSE for 2 hours, 4 hours, and 6 hours, followed by 16-20 hours of recovery in fresh media. We designed the experiments described in **Fig. 1** with three sets: siSCAL1 (-), siSCAL1 (+), and siControl (+) BEAS-2B cells.

### Intracellular ROS estimation

The BEAS-2B (5 x 10^7^ cells/ml) cells were incubated with a membrane permeable non-fluorescent probe, 10 µM of 2’, 7’-dichlorodihydrofluorescein diacetate (H2-DCFDA; Sigma-Aldrich) at 37^°^C for 30 mins in a dark humidified chamber with 5% CO_2_. The amount of intracellular ROS was estimated by measuring the fluorescence intensity of the oxidized product, 2’, 7’-dichlorodihydrofluorescein (DCF) at excitation and emission wavelengths of 488 nm and 522 nm, respectively, in a fluorescence spectrometer. Furthermore, fluorescent images were captured in the 488 nm filter using the 40x objective of an inverted microscope (Olympus IX71).

### Preparation of cDNA from RNA by RT-PCR

Total RNA was isolated by TRIzol® (Invitrogen, Carlsbad, CA), following the protocol by Chomczynski P. et al. [76]. The total RNA was quantified in NanoDrop 1000 (ThermoFisher Scientific). About 1µg of total RNA was reverse transcribed to cDNA with the iScript cDNA Synthesis kit (Bio-Rad Laboratories, Hercules, CA; #1708891) following the manufacturer’s protocol.

### Quantitative Real-Time PCR

We performed real-time qPCR reactions in Applied Biosystems OneStep^TM^ thermocycler with QuantStudio 5 software. We quantified SCAL1 expression in BEAS-2B cells at 20% CSE exposure for 2 hours, 4 hours, and 6 hours, followed by 16-20 hours of recovery time. The specificities of the amplification were analyzed and confirmed from the melting curve of the reaction. We estimated the relative mRNA abundance to the external calibrator gene β actin by the *Livak* method (2-ΔΔCt). The results were expressed as fold change compared to the external calibrator. Specific primers sequences for SCAL1 (forward: 5’-ACCAGCTGTCCCTCA GTGTTCT-3’, 5’-AGGCCTTTATCCTCGGGTTGCCT-3’: reverse) and β-actin (forward: 5’-GCGGGAAATCGTGCGTGACATT, 5’-GATGGAGTTGAAGGTAGTTTCGTG-3’: reverse) were taken from Thai et al. [10]. The primer sequences were purchased from Integrated DNA Technologies (USA). Each reaction was prepared in biological and technical triplicates for all the treatment conditions.

### Immunofluorescence Imaging

BEAS-2B cells were cultured on poly-L-lysine coated coverslips in sterile cell culture grade 35 mm dishes. After the usual 20% CSE treatment and siSCAL1 transfection procedure, the culture plates were kept on ice, and the medium was aspirated. We followed Abcam’s Immunocytochemistry and immunofluorescence staining protocol (https://www.abcam.com/protocols/immunocytochemistry-immunofluorescence-protocol). Following this, the images were captured by an inverted fluorescence microscope (Olympus IX71) with green, red, and blue filters. The fluorescence signal was quantified using ImageJ software (https://imagej.nih.gov/ij/), and the corrected total cell fluorescence (CTCF) was calculated using the following formula [77]:

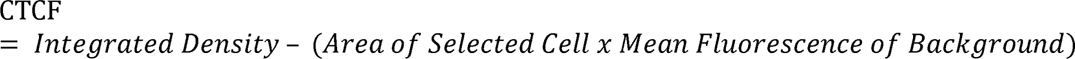

### Western Blot Analysis

About 40 µg of total protein was mixed 1:1 v/v with 2x sample buffer (20% glycerol, 4% SDS, 10% β-mercaptoethanol, 0.05% Bromophenol blue, and 125mM Tris-HCl of pH 6.8) and following the standard protocol. The primary antibodies, like anti-NRF2 (#12721T, Cell Signaling Technology®), anti-P53 (#9282T, Cell Signaling Technology®), anti-CD44 (#3578S, Cell Signaling Technology®), anti-CD133 (5860S, Cell Signaling Technology®), anti-TCTP (#5128T, Cell Signaling Technology®), anti-APAF1 (#4452, Cell Signaling Technology®), anti-Merlin (#12888, Cell Signaling Technology®), anti-PLK1 (#4535, Cell Signaling Technology®), anti-ALDH1A1 (#12035, Cell Signaling Technology®), anti-BCL2 (#4223, Cell Signaling Technology®), anti-CASP-3 (#14220, Cell Signaling Technology®), and anti-CytoC (#4272, Cell Signaling Technology®) at a dilution of 1:1000 in 5% non-fat dry milk in 1X TBS overnight at 4^◦^C in a platform rotary shaker. The membranes were washed thrice with 1xTBST buffer for 5mins each and then incubated for 2 hours with alkaline phosphatase-conjugated anti-rabbit IgG (Abcam; #ab97048) at dilutions of 1:10000 in 5% milk TBST. Following the standard protocol, the immunoreactive bands were developed by incubating the membranes with Alkaline Phosphatase Chromogen (BCIP/TNBT; Abcam, #ab7413). Images were taken in ChemiDoc^TM^ XRS+ System (Bio-Rad) with exposure varying between 30 and 90 secs. Equal loading was confirmed with primary antibody anti-α tubulin (#2144, Cell Signaling Technology®) at a dilution of 1:1000 and alkaline phosphatase-conjugated anti-mouse IgG (Abcam, #ab97020) at a dilution of 1:10000, following the same steps mentioned above. Densitometry analysis was performed using ImageJ software (https://imagej.nih.gov/ij/).

### Soft Agar colony formation assay

The tumorigenic potency of siSCAL1 (+) and siSCAL1 (-) BEAS-2B cells on 20% CSE exposure was determined using the soft agar colony formation assay according to the protocol of Borowicz et al. [78] with modifications. The bottom agar was prepared by pre-warming 50 mL of 1% agar in 2x complete media (1:1 v/v) at 42°C in a hot water bath, then transferring 2 mL of the pre-warmed mixture into clear flat-bottomed sterile 6-well plates (Corning®). The plate was chilled at 4°C to solidify the bottom agar and prevent cell adhesion. Single-cell suspensions of the cells were prepared using a 0.25% trypsin-EDTA solution (Gibco, MA, USA) and adjusted to the targeted cell density of 10,000 cells/mL by diluting with complete media. The cells were then mixed into 0.8% agar at a 1:1 ratio to obtain a final concentration of 0.4% agar containing 5000-6000 cells/mL. A total of 6 mL of the 0.4% agar-containing cells was layered onto the 0.5% bottom agar in each well of the 6-well plates, which were then chilled at 4°C for 20 min to solidify the cell agar. Complete serum-free media (with supplements) was added to each well (2 mL per well), and the cells were cultured at 37 °C in 5% CO2. The assays were performed in triplicates with 20% CSE exposure for 4 and 6 hours daily, followed by 16-18 hours of recovery in fresh, complete, serum-free media on siSCAL1-treated and siSCAL1-untreated BEAS-2B cells. The medium was changed every 3^rd^ day until the 11th and 21st days. The cells were not stained for visualizations. The number of spheroid colonies was assessed on the 11^th^ and 21^st^ day from baseline (Day 0; cell seeding) for siSCAL1-treated and siSCAL1-untreated BEAS-2B cells under CSE exposure as earlier defined.

### Interactome Analysis

We performed an interactome analysis in STRING v12 (http://string-db.org) [79, 80], including known and predicted protein-protein interactions. The interactome was expanded to gain more interactors, with a required confidence score > 0.4 as the cut-off value.

### Statistical Analysis

All experiments were performed in triplicate and repeated three times for independent cell passages. Data were checked for normality using the Shapiro-Wilk test. Normally distributed variables were summarized as mean ± standard deviation (SD), while non-normally distributed variables were presented as median. Mean±SEM measurements were taken for multiple instances for graph plotting. Data were analyzed using one-way analysis of variance (ANOVA) followed by post hoc analysis using the Bonferroni method. Statistical significance was set at *p*<0.05*. R Version 3.4.2 was used for data analysis [81].

## Acknowledgments

This study was supported by the Department of Science and Technology (DST), Government of India—Promotion of University Research and Scientific Excellence (DST-PURSE) provided to the University of Calcutta. We acknowledge the University Grants Commission, Government of India, for providing the UGC UPE-II Grant: Focus area: Modern Biology Group C1: Investigating mechanistic approaches for delineating proliferative diseases (Grant No. not applicable) to conduct this study. The funding agency had no role in the design, execution, and publication of the study. No financial assistance is provided for the publication of the study findings. We also express our gratitude to Dr. Anurag Agarwal, Sr. Scientist and Director of Translational Research in Lung Disease at the Institute of Genomics and Integrative Biology (IGIB), New Delhi, India, for generously gifting us the BEAS-2B cell line, and to Dr. Somsubhra Nath, Assistant Professor, Institute of Health Sciences, Presidency University, Kolkata, West Bengal, India for gifting us the scrambled siRNA controls and antibodies to conduct the study. We also thank Dr. Alok Sil, Professor, Department of Microbiology, Dr. Sagartirtha Sarkar, Department of Zoology, and Dr. Aniruddha Mukherjee, Dr. Punarbasu Chaudhuri and Dr. Pritha Bhattacharjee of the Department of Environmental Sciences, University of Calcutta, Kolkata for providing access to the respective departmental and laboratory instrument for the conduct of the study.

## Authorship Contributions

DS and MS: **Conceptualization**. DS and MS: **Methodology**. DS, SB, and MS: **Investigation**. DS and MS: **Project Administration**. DS, SB, and MS: **Resources**. DS: **Formal analysis**. DS and SB: **Validation**. DS: **Writing-Original draft**. DS, SB, and MS: **Writing-Reviewing and editing**. MS: **Supervision.**

All the authors have reviewed the manuscript and have agreed to publication.

